# SIS-seq, a molecular ‘time machine’, connects single cell fate with gene programs

**DOI:** 10.1101/403113

**Authors:** Luyi Tian, Jaring Schreuder, Daniela Zalcenstein, Jessica Tran, Nikolce Kocovski, Shian Su, Peter Diakumis, Melanie Bahlo, Toby Sargeant, Phillip D. Hodgkin, Matthew E. Ritchie, Shalin H. Naik

## Abstract

Conventional single cell RNA-seq methods are destructive, such that a given cell cannot also then be tested for fate and function, without a time machine. Here, we develop a clonal method *SIS-seq*, whereby single cells are allowed to divide, and progeny cells are assayed separately in *SIS*ter conditions; some for fate, others by RNA-*seq*. By cross-correlating progenitor gene expression with mature cell fate within a clone, and doing this for many clones, we can identify the earliest gene expression signatures of dendritic cell subset development. SIS-seq could be used to study other populations harboring clonal heterogeneity, including stem, reprogrammed and cancer cells to reveal the transcriptional origins of fate decisions.

## Main text

Single cell analyses including flow cytometry, microscopy, colony assays, clonal lineage tracing, and most recently single cell genomics methods, have revolutionised our understanding of biological systems and their heterogeneity^1^. As demonstrated by clonal assays, haematopoietic stem and progenitor cells (HSPCs) are particularly heterogeneous in their fate^2^, and the molecular programs governing these are gradually being characterised^3-7^.

One challenge for connecting transcriptional signatures with functional heterogeneity is that these properties can rarely be measured on the same single cell *i.e*. single cell RNA-seq is destructive, so the same cell cannot then be tested for fate and, *vice versa*, a single cell tested for fate divides and differentiates such that the founder cell cannot be tested for its molecular profile. A time machine could conceivably allow one to first ascertain one feature of a given cell, then go back in time and re-test the same cell for the other feature, allowing cross-comparison of the two. This is reminiscent of the challenge in quantum mechanics where a particle’s momentum and location cannot be simultaneously known (the popular interpretation of the ‘Heisenberg uncertainty principle’).

For single cell biology, we reasoned that one solution to this challenge could be if single HSPCs were allowed to undergo limited clonal division such that resulting clones could be split, and siblings used separately in assays of both single cell RNA-seq and fate. In this way, siblings would be surrogates of the founder for later cross-correlation of fate heterogeneity with molecular heterogeneity. We term this approach of testing split clones in different *SIS*ter conditions for RNA-*seq,* or fate, as *SIS-seq*.

For this approach to work, several important conditions need to be met; a) that daughters derived from single stem or progenitor cells are highly concordant in their fate; b) that within an HSPC population there is fate heterogeneity allowing differences to be resolved, and c) that this program is reflected in measurable molecular features.

We chose dendritic cell (DC) development as a suitable test of SIS-seq. DCs are a family of immune cells including conventional DC type 1 (cDC1), cDC type 2 (cDC2) and plasmacytoid DC (pDC), important for the detection of perturbed immune homeostasis, as well as immunity against pathogens, cancer and self-antigens^8^. The prevailing model suggests that DCs can be generated from HSPCs via Flt3-expressing progenitor cells^9,10^, which transition to a common DC progenitor (CDP)^11,12^, and on to dedicated progenitors^13-15^ for the individual DC subtypes. Relevant to SIS-seq is that DCs can be generated *in vitro*^16^ for ease of assessment of multiple clones, and be derived from single HSPCs^11^. Moreover, cellular barcoding and clone-splitting experiments *in vivo*^17^ and *in vitro*^18^ demonstrated that DC lineage bias, and also individual DC subtype bias, is heterogeneous amongst HSPCs, but restricted within a given clone. These properties fit the first two conditions for SIS-seq to be feasible. Moreoever, known molecular regulators of DC subtype fate can be identified within single cells of DC progenitors^14^, thus fitting the third condition. However, as most of these molecular regulators that have been identified to act at downstream progenitor stages of DC development, closer to the mature cell^19^, there remained an opportunity to identify novel earlier regulators in less-differentiated HSPCs.

SIS-seq was used to identify the early transcriptional origins of DC fate from HSPCs, as follows: Single Sca1^+^ ckit^+^ HSPCs were isolated from BM of ubiquitin C-GFP mice, which allowed their tracking, and cultured amongst a pool of non-GFP BM filler cells from C57BL/6 mice in medium supplemented with Flt3 ligand^11^. Clones were split after 2.5 days, which allowed sufficient progenitor cell expansion for clone-splitting, but which was prior to DC differentiation^11^. At this time, wells were examined by microscopy for GFP^+^ clones, and those with 10 or more cells were split into three equal parts to yield duplicate wells for further culture to test fate at day 8, and the remaining third for single cell, or small cell number RNA-seq using CEL-Seq^20^ on the less differentiated cells to provide a transcriptional ‘snapshot’ of the clone prior to differentiation.

We examined the fate of 105 clones in duplicate split culture conditions using FACS. Most clones were conserved in their lineage bias between sister wells. Four example clones are shown in Figure 1b, with different indicated fate biases. To better visualise fate conservation for all clones, we used ternary plots (Figure 1c), where a sample’s cellular composition dictates its position. For example, where a sample contained only pDCs, it would be at the apex of the plot marked pDC. If the sample contained equal numbers of the three DC subtypes, then it would be placed in the centre.

**Figure 1.**
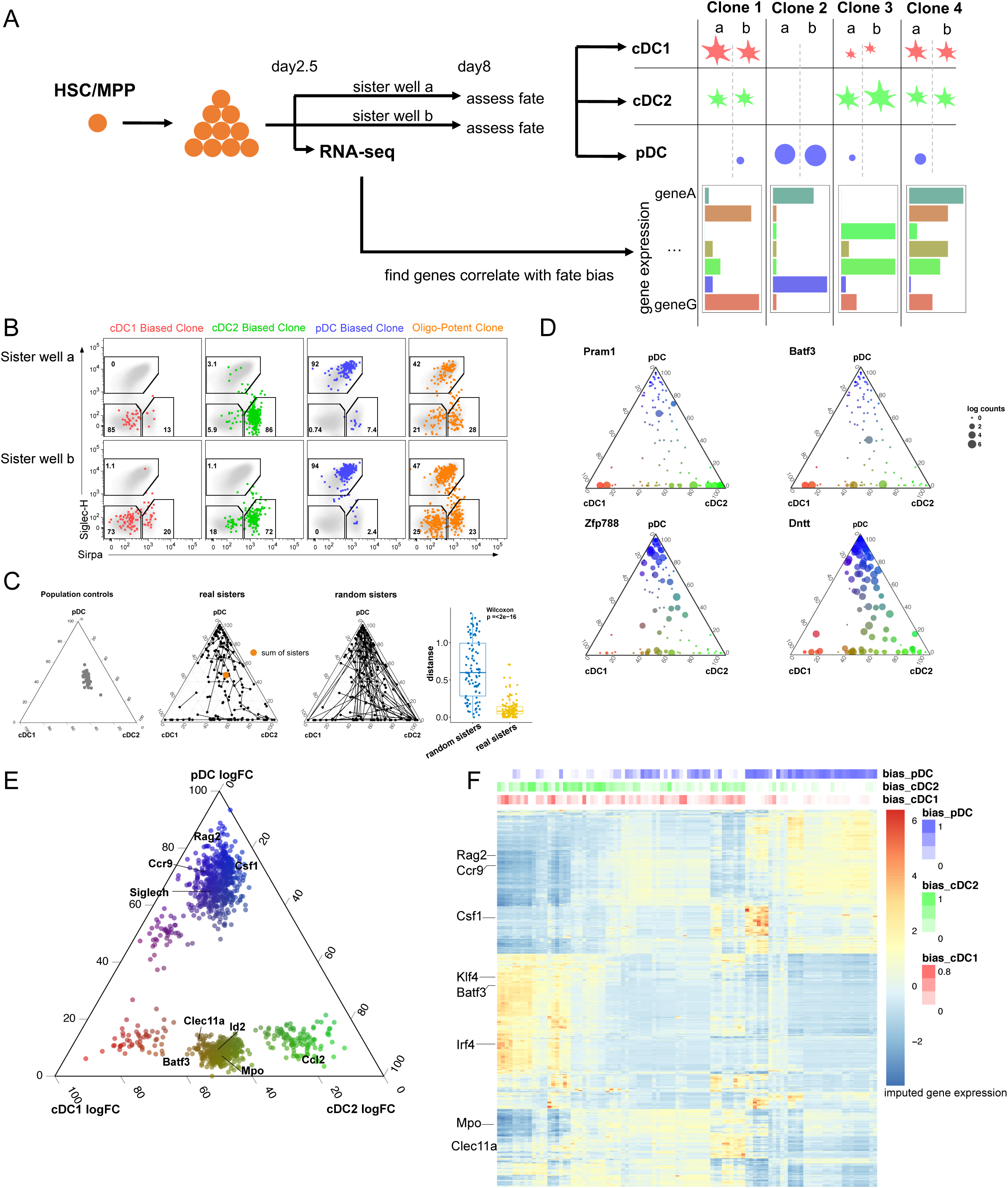
SIS-seq captures gene programs in DC subset development. (A) The conceptual framework of SIS-seq. After a small pre-expansion of single HSPCs, clones are subdivided into three parts at day 2.5; one for RNA-seq, and other two further cultured for fate assays in ‘sister’ wells. Fate outcomes are then measured by flow cytometry, and computationally correlated with gene expression of sibling cells from day 2.5. (B) Example FACS plots showing conserved fate bias between siblings of pre-expanded HSPC clones in DC culture assays. (C) Fate bias in ternary plots of internal control samples (left), clonally-derived DCs where sister clones are connected (middle ternary plot) including the sum of all clones (orange dot), or where sister wells are randomly connected (right ternary plot). The distance between real sisters and random sisters summarized in the adjoining box plot for all clones. (D) Ternary plots show correlation between gene expression and fate bias for both novel and known marker genes. Each clone was averaged and placed as a single dot, then the size of that dot altered according to the expression of the indicated gene. (E) Ternary plots of scaled log fold change of fate bias for significant genes, with genes known to be associated with DC fate marked. (F) Clustered gene expression heatmap of significant genes across 98 clones, with the fate bias of each clone towards pDC, cDC1 and cDC2 layered on top.

We first assessed the internal non-GFP population controls to establish whether any well exhibited any fate bias. As seen in Figure 1c, left ternary plot, this was not the case. When we assessed the fate bias of GFP^+^ cells for all clones within those same wells, and connected duplicate wells with a line to visualise their fate relationships, we observed that clones were highly heterogeneous in their fate, but fate was similar for those clones split into sister wells (short connecting lines). Moreoever, when the individual clones were summed together, they recapitulated the cellular composition of the population controls (Figure 1c, middle ternary plot, orange dot). This contrasted when those same wells were randomly connected to another clone, which exhibited long lines (Figure 1c, right ternary plot). By using Euclidean distance between the fate of sisters, which is a statistical readout for similarity in fate, we demonstrated real sister clones were more conserved in their fate (small distance) than random sisters (large distance) (Figure 1c, right panel). Therefore, while HSPC clones were heterogeneous in their fate, clones in sister conditions were highly conserved. Thus, the aforementioned condition of conserved fate for SIS-seq to be useful was met for most clones (98/105). Those clones not conserved in fate may be of biological interest, but were excluded from further analyses to focus on the initial goals of *SIS-seq*.

Using clones with conserved fate, we sought to identify those genes expressed in their less differentiated siblings that correlated with each clone’s final mature DC fate, using linear regression analysis (see Methods). As expected, we identified genes known to be associated with DC subtype-specific fate such as *Batf3* in cDC fate^21^, and *Dntt* with pDC fate^22^, however the analyses also revealed novel genes such as *Pram1* and *Zfp788* (Fig 1d). More than 1000 genes were determined to be associated with individual or dual DC subtype output as can be seen in a ternary plot (Figure 1e, Supplementary Table 1). Here, the position of the gene is proportional to its enrichment within a given fate bias. This data was also visualised as a clustered heatmap (Figure 1f) where clustering was only based on expression of genes selected by SIS-seq, and fate was layered on top afterwards. As observed in Figure 1f, fate segregated into clusters generated by RNA-seq. The genes were further filtered by the glmnet algorithm, and ribosomal and mitochondrial genes eliminated, resulted 613 genes as our final list (Supplementary Table 2).Thus, only by assessing many clones, and being able to link gene expression with fate of clones using SIS-seq, were the transcriptional correlates of early DC fate observed.

To understand whether other approaches and datasets could have identified similar gene expression correlates of fate, we compared them to those using SIS-seq. First, we asked whether the most highly variable genes in the same clonal data (i.e. in the absence of fate information) would have derived a similar set of genes. However, only 40 out of those top 600 variable genes was shared with the 613 identified with SIS-seq, and did not include several known factors in DC development, including those sparsely expressed within the clonal data such as *Batf3* (data not shown). This unsurprising but important result confirmed that the supervised analyses for fate in SIS-seq yields a unique gene set.

We also compared SIS-seq to approaches where candidates were identified based on differential expression between: a) mature DC subtypes or, b) CDPs and other HSPC populations from the Immgen database^23^, or c) highly variant from single cell RNA-seq data of CDPs from a published dataset^14^ (Figure 2a). We assessed the overlap of genes identified using SIS-seq and those identified by these alternative approaches (Figure 2b) and 383/613 were identified using SIS-seq and not other approaches.

**Figure 2.**
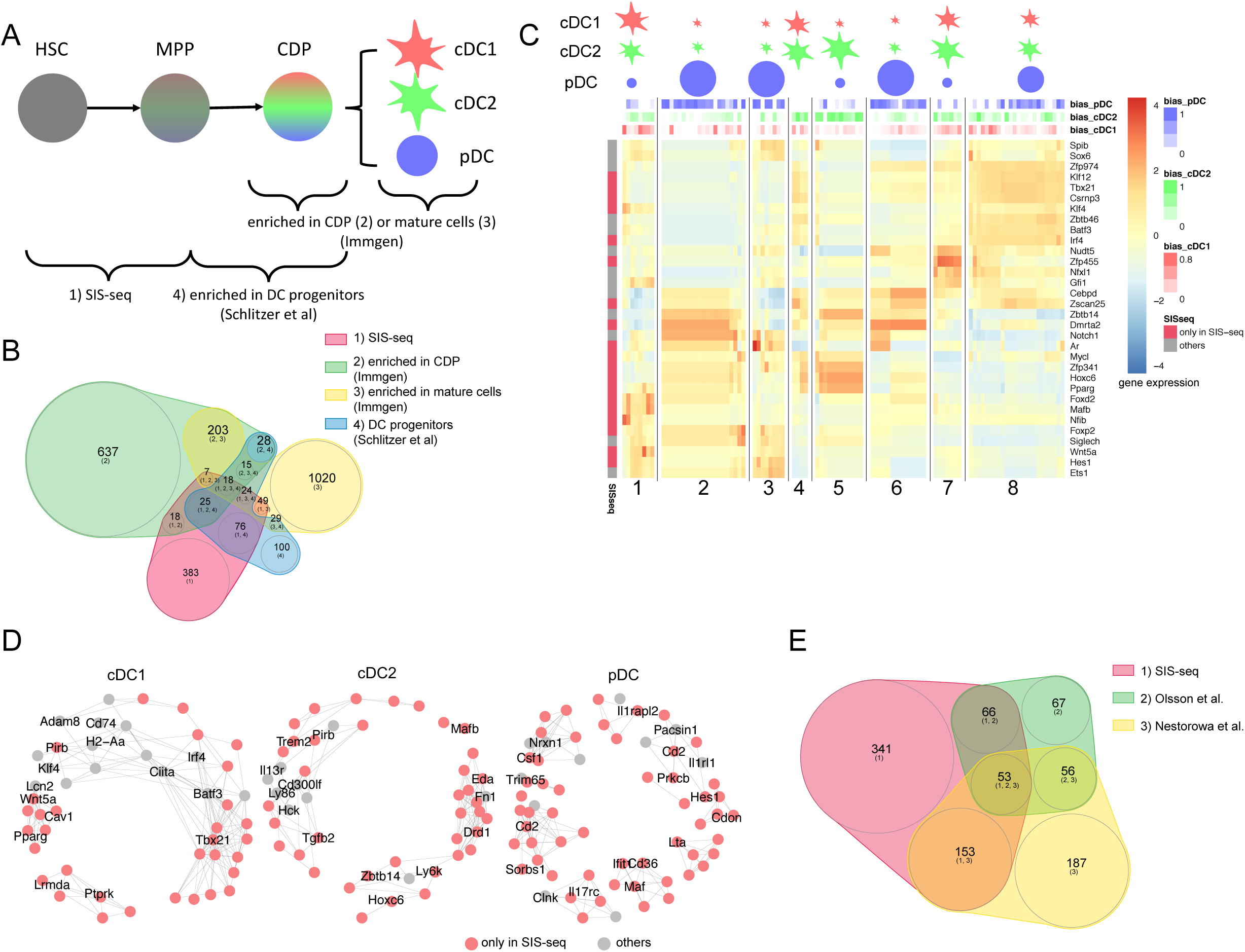
SIS-seq highlights unique gene expression signatures in DC development. (A) Genes derived using SIS-seq compared with other gene lists, including those highly expressed in CDPs or mature DCs using Immgen; and variable genes expressed in single DC progenitors. These data sets correspond to different stages of DC development as indicated such that SIS-seq identifies the earliest gene expression correlates. (B) The overlap among different gene sets in (A) is highlighted in the Venn plot. (C) The gene expression of transcription factors identified with SIS-seq, which include known marker genes in DC development (grey annotation in y column) as well as novel genes (pink annotation in y column). Clones are portioned into 8 clusters based on gene expression, with different lineage bias patterns highlighted on top. (D) Gene co-expression network of regulatory genes that correlated with fate bias. Genes are coloured based on the overlap in Figure 2B, where genes in grey could be identified from prior data, and those in pink are novel. (E) The overlap between SIS-seq identified genes to highly variable genes in other HSPC datasets.

Transcription factors and epigenetic regulators are good candidates for genes important in fate determination. Those identified by SIS-seq are shown in Figure 2c, where 19/32 were identified only by SIS-seq and not the other techniques. This list included several known genes involved in fate^24^ determination including *Batf3, Irf4* and *Zbtb46*, but also novel genes including *Hoxc6, Dmrta* and *Mafb* (annotated by row in Figure 2c, based on Figure 2b). Thus, SIS-seq identified known genes but also revealed novel gene expression programs that correlate with fate. Some genes known to be involved in the development of specific DC subtypes were not identified, such as *Tcf4*. This was not surprising considering SIS-seq identifies the earliest gene expression programs of fate bias, not those differentially expressed late in DC development, such as *Tcf4*^25^. The coexpression network of regulatory genes that correlated with fate bias of the three DC subtypes was created (Figure 2d) and we found the genes identified only by SIS-seq were coexpressed with known marker genes.

Notably, there was an imperfect correlation between transcriptional clusters and fate bias e.g. clones within clusters 2, 3, 6 and 8 were pDC-biased but expressed different gene signatures. There could be three possibilities for this: a) there were different molecular routes to the same cell type, and there is precedence for this in DC development^26^; b) the clusters represent different points of the same developmental trajectory, and the snapshot of our clones captured these different stages; and c) there is heterogeneity within our flow cytometric-sorted populations of DCs and the clusters may represent different clonal routes to these putative subpopulations. Recent evidence suggesting Siglec-H^+^ cells can be further separated by CCR9 expression into cDC progenitors and *bona fide* pDCs^18,27^, so this might be an example of c) as pDCs were not sorted based on CCR9 in these experiments. That is, Cluster 8 may represent genes that actually correlate with cDC bias, and where cells phenotypically ascribed as pDCs were in fact CCR9^−^ cDC progenitors. In contrast, clusters 2, 3 and 6 appear similar, and may better reflect pDC differentiation programs, albeit at different stages or pathways.

Lastly, we sought to understand whether recent scRNA-seq of HSPCs might have identified similar genes, considering our starting population was HSPCs rather than the downstream progenitors of Figure 2 a-d. Using two published datasets^5,6^, we determined a number of highly variable genes based on a statistical cut-off that gave a similar number of genes identified using SIS-seq. Only 55% were shared (Fig. 2e), indicating that scRNA-seq alone does not always identify those genes that correlate with a particular fate.

In summary, we demonstrate SIS-seq as a simple and powerful approach to uncover many subtle differences in gene expression that correlate with fate, which would otherwise be difficult to determine with standard study designs and scRNA-seq, or other single cell approaches. We envisage that SIS-seq, which approximates a molecular ‘time machine’, can be used to discover the features of rare subsets of cells based on their fate or function, and will be of particular interest in stem cell research, cancer and reprogramming. This approach could also incorporate any other ‘omics measurements, such as ATAC-seq, that can be assessed as single cells or using small cell numbers. As most multi-omics methods strive to separate molecular features on the same single same cell^28^, which can be difficult to achieve, SIS-seq could provide a more practical alternative using separate but clonally related cells *i.e*. clonal multi-omics.

SIS-seq could be scaled-up and deployed with higher resolution, for example using single cell transcriptomics, and an expressed lineage tracing barcode^29^. Barcoded progenitors could be pooled in experiments, and one third of cells assessed by scRNA-seq and endogenous barcode expression at the time of clone-splitting, followed by assessement of barcode distribution in the cell types derived from the other two thirds of cells cultured until the end point of the experiment. By systematically linking fate with molecular heterogeneity of the undifferentiated cell using SIS-seq, we envisage that refined maps of haematopoiesis and other developmental processes are achieveable.

## Methods

### Mice

All mice were bred and maintained under specific pathogen-free conditions at WEHI, according to institutional guidelines. Bone marrow cells from C57BL/6 (CD45.2) were used as filler cells, and C57BL/6-TG (UBC-GFP) 30scha/j as a source of GFP^+^ single cells. Mice were between 8-16 weeks at the time of sacrifice.

### Flt3 ligand cultures of DCs

Cultures were set up as described^16,30^. Briefly, bone marrow cells were extracted from C57BL/6 mice and cultured at a density of 1.5 × 10^6^ in 96-round bottom plates with DC culture medium and 200 ng/ml FLT3L obtained from Dr Jian-Guo Zhang (WEHI). To generate conditioned medium that can sustain the growth and differentiation of isolated DC progenitors, medium derived from FLT3L cultures after 3.5 days was centrifuged to remove cells, then passed through a 0.22-μm filter, and frozen aliquots stored at −80°C, as described previously. This medium is referred to as DC-conditioned medium (DC-CM).

### Isolation and sorting of single GFP^+^ MPP or LSK cells

The isolation and sorting of GFP^+^ progenitors was performed as described^11^. Briefly, bone marrow from C57BL/6-TG (UBC-GFP) was extracted and cells were enriched for ckit (2B8 conjugated to APC) via magnetic activated cell sorting (MACS) positive selection. The ckit enriched fraction was subsequently stained for Sca-1 (E13 161-7conjugated to Alexa Fluor 594 or Alexa Fluor 680), IL-7R (A7R34-2.2 conjugated to biotin with streptavidin-PECy7) and FLT3 (A2F10 conjugated to Phycoerythrin). Cells with positive expression of GFP, ckit, sca-1 and FLT3, while negative for IL-7R, were deemed MPPs and sorted into FLT3L-DC “filler” cultures as single cells, with 100-cell controls in duplicate. Cells with positive expression of GFP, cKit and Sca-1 were deemed Sca1^+^cKit^+^ (SKs). Cells were sorted using the BD Influx, BD FacsARIA W and BD FacsAria L. Single SKs from UBC-GFP mice were deposited in 96 well round bottomed wells pre-seeded with 1.5 × 10^6^ total C57BL/6 BM cells (filler cells) cultured in DC-CM with an additional 200 ng/mL FLT3L at 37 °C and 5% CO_2_ for 2.5 days.

### Day 2.5 split and fluorescence imaging

On day 2.5 after culture, each 200 μl cultured well was mixed well, then 100 µL of the contents transferred manually to a separate new well, and given approximately 30-60 minutes to settle. The 96-well plates containing sister wells derived from each single UBC-GFP LSK clone were imaged on a Nikon TiE live cell imaging system. Cells were maintained at 37 °C in a humidified atmosphere and 5% CO_2_. Images were acquired in DIC and fluorescence using a 4x 0.13NA plan neofluor objective, and GFP filter (FITC, ex 465-495, DM 505, em 515-555). A Z-stack of 150 μm, with interval of 50um was performed and, the best focus plane was automatically selected for analysis using an offline Metamorph software (v 7.8.2.0) licence and method for automatic segmentation and cell count. Cell count script parameters, region size and diameter, were changed per experiment, on the basis of visual inspection of the 100 single cell controls. After imaging, cell cultures were incubated in 37 °C and at 5% CO_2_ for 4.5 days.

### Day 8 imaging and FACS analysis of DC clones

After a further culture period of 6 days (total of 8 days from the original progenitor deposition), 96-well plates were imaged on the Nikon TiE live cell imaging system as described above. Wells with clones of greater than 10 cells were selected for flow cytometric analysis. Cells were labelled with antibodies to CD11c (N418-APC), Sirpα (P84 conjugated to Alexa flour 594 or 680) and Siglec-H (551 conjugated to PE). Flow cytometry was performed using a BD FACS Canto II (BD Biosciences) and BD Fortessa X20 (BD Biosciences). DCs were labelled with forward and side scatter profiles, with gating to remove doublets. To select for DCs, cells were gated based on intermediate to high expression of CD11c-APC, with subtypes gated for Sirpα and Siglec-H. DC that had negative expression of Sirpα and Siglec-H were cDC1s, those that were Sirpα^+^ were cDC2s, and pDCs were classified expression of Siglec-H. Data were analysed using FlowJo software (v 7.6.5), Microsoft Office Excel 2007 and Graphpad Prism (v 6.01).

### RNA-seq library preparation and data preprocessing

Depending on cell abundance 1-10 cells per clone were FACS sorted for RNA sequencing. Transcriptome libraries were generated according to the CEL-Seq protocol^20^ with the exception that all column purifications were replaced by magnetic bead purifications using AMPure XP beads (Beckman Coulter A63880). Libraries were sequenced on the Illumina NextSeq500 and MiSeq sequencer using a 75-cycle kit. Raw data are available in GEO under accession number GSE119097.

The data was first demultiplexed by the cell barcode sequence and then aligned and counted using Rsubread (1.5.0)^31^ and featureCounts (1.5.0)^32^ with mm10 genome annotation. Quality control was performed using scPipe^33^ to remove low quality samples. The resulting gene count matrix was used as input for downstream analysis, and normalized using the same method as in Velten *et al*^3^ for further analysis.

### Examining lineage bias with gene expression data

The relative proportion of cells within each gate of the DC subtypes was determined by FACS for each well, and then the statistical distance between sister wells calculated, and used together with the total cell count as the input of Mahalanobis distance for outlier detection. Random permutation on sisters described in Figure 1b were used to get a null distribution and a 0.05 *p*-value cut-off was used to exclude sisters with different lineage output, leaving us with 98 clones. We used edgeR (3.20.8)^34^ to model the correlation between lineage bias and gene expression, with lineage bias and batch as covariates in the generalized linear model. A *p*-value cut-off of 0.01 was used to select genes that correlate with lineage bias, which gave us 1071 genes (Figure 1e, f). The fold changes (FC) for each genes were scaled to 0-1 and converted to percentage and ploted in Figure 1e. Similar to STEMNET, we then applied a generalized linear model with regularization on the selected genes using glmnet^35^ to get the final SIS-seq prioritised gene list with 613 genes (Figure 2b). The parameter for regularization was chosen by 10-fold cross validation. The gene list was further reduced by selecting known marker genes and regulatory genes such as transcription factors and epigenetic modifiers using GO terms (Figure 2c). The gene expression matrix was imputed using MAGIC (0.1.0) ^36^ before plotting heatmaps using pheatmap (https://github.com/raivokolde/pheatmap). The cubed pairwise Pearson correlation of SIS-seq genes in Figure 2b was computed in order to reduce noise in correlation structure, and genes with correlations higher than 0.15 were selected. Correlations with lineage bias were also calculated and genes with correlation higher than 0.15 were kept in the co-expression network (Figure 2d).

### Comparisons to public datasets

For the DC dataset comparison in Figure 2b, differentially expressed gene lists were downloaded from the Immgen website (immgen.org)^23^. The genes enriched in CDP were derived by comparing CDPs to other progenitors, including GMPs, CMPs, CLPs and MDPs. The genes that were enriched in mature cDC1, cDC2 and pDC were obtained by comparing against the other two subtypes. The gene count matrix from Schlitzer et al. was downloaded from GSE60781. The informative genes were generated using the methods described in the scran pipeline^37^, which is to filter the highly variable genes and then perform gene-gene correlation analysis, keeping the correlated genes pairs (FDR<0.05). For the HSPC dataset comparison in Figure 2e, the two public HSPC datasets were downloaded from GSE81682 and GSE70245. The “HSC”, “LT-HSC” in GSE81682 and “LSK” in GSE70245 were selected to represent HSPCs. The informative genes were selected in a similar way as for Schlitzer *et al*.^14^, which was to pick highly variable genes and keep correlated gene pairs. All venn diagrams were ploted using the nvenn package^38^

